# Changes in length-at-first return of a sea trout (*Salmo trutta*) population in northern France

**DOI:** 10.1101/2023.11.21.568009

**Authors:** Quentin Josset, Laurent Beaulaton, Atso Romakkaniemi, Marie Nevoux

## Abstract

The resilience of sea trout populations is increasingly concerning, with evidence of major demographic changes in some populations. Based on trapping data and related scale collection, we analysed long-term changes in body length of a sea trout population in the Bresle River, France. From 1984-2022, the length of first-time returning individuals decreased by 1.73 mm.year^-1^ (SD = 0.08), which resulted in the loss of *c.* 12.3% of mean body length. This decrease results from a decrease in the age at first return, with a gradual loss of the oldest individuals and an increase in the proportion of the youngest. Timing of the return migration advanced drastically, although shorter sea sojourn had little influence on body length. We found little evidence of a decrease in length-at-age, to the exception of the oldest age class, suggesting that growth conditions at sea might not have deteriorated greatly during the study period.

## Introduction

The environmental conditions that individuals experience, in conjunction with their genotype and epigenetic mechanisms (Ferguson et al., 2019), are susceptible to influence the expression of their phenotype and their life history trajectories (Walsh and Reznick, 2011). Migration decision, survival, growth, age at maturation or fecundity can thus be influenced by a series of intrinsic and extrinsic factors, such as environmental conditions (Crozier and Hutchings, 2014; Satterthwaite et al., 2012), genotype (McKinney et al., 2015), inter- and intra-specific competition (Helle et al., 2007), predator-prey relationships (Ohlberger et al., 2019), and anthropogenic activities (Hard et al., 2008; Law, 2000).

Recent environmental changes attributed to climate change and anthropogenic activities are assumed to be responsible for the widespread decline in salmonid abundance and distribution (Forseth et al., 2017; Kendall and Quinn, 2011; Kennedy and Crozier, 2010; Perrier et al., 2013). Ongoing changes, such as an increase in sea temperature, ocean acidification and stratification, rising sea levels, and eutrophication, are likely to influence marine trophic conditions (Möllmann and Diekmann, 2012). These changes could greatly modify the quality, quantity, and availability of trophic resources in the ocean (Bartolino et al., 2014; Brown et al., 2010; Niiranen et al., 2013) and thus impact growth opportunities for salmonid populations at sea.

Reduced growth opportunities can influence individuals’ life history trajectories and fitness greatly by decreasing survival, fecundity, and competitive abilities (Perez and Munch, 2010; Quinn et al., 2016) or by delaying the age at maturation (Ishida et al., 1993). These changes can cause large variations in the length and age structure of populations. Such changes in population composition have been recorded for many salmon populations around the world (Bal et al., 2017; Jeffrey et al., 2017; Lewis et al., 2015; Morita and Fukuwaka, 2007; Ohlberger et al., 2018; Oke et al., 2020), but also sea trout (Milner et al., 2017), and could indicate a widespread decrease in marine growth and survival. In this context, identifying the ecological mechanisms that most likely influence these changes is crucial to better understand and predict population dynamics. This knowledge is also necessary to provide recommendations for effective management actions and to alleviate or compensate for pressures on sea trout populations when and where possible.

The anadromous form of the brown trout (*Salmo trutta,* L.) (hereafter, “sea trout”) (ICES, 2020) exhibits an extreme diversity of life histories across its wide distribution range (Jonsson and L’Abée-Lund, 1993), which makes it an appropriate model species to study variations in life histories in a changing environment. The age at seaward migration (*i.e.* smolt stage) ranges from 1-3 years in France to 5-7 years in northern Norway (Nevoux et al., 2019). First reproduction occurs after 0-2 or 2-4 years at sea in populations in the English Channel (Richard, 1981) and the Baltic Sea (Järvi, 1940), respectively.

In recent decades, alarming collapses in population abundance, as well as altered population structure, have been recorded in Europe, such as in the Burrishoole River, Ireland (Gargan et al., 2006), the Ewe River, Scotland (Butler and Walker, 2006), and the Vistula River, Poland (Dȩbowski, 2018). Damming of rivers forbidding the access to spawning sites, overfishing, or the development of offshore salmon aquaculture leading to sea lice epizootics, have been pointed as major factors of decline for some of these populations. Consequently, it is necessary to assess and understand possible changes in key life history traits, such as length at spawning, age at maturation, and length-at-age, which drive the dynamics of European sea trout populations.

In this study, we quantified the influence of key life history and phenological variables on the length of sea trout returning to the Bresle River, France, using 39 years of data from long-term population monitoring. Based on an intensive trapping protocol throughout the migration season, we obtained individual data on length, migration timing, and age, as inferred from scale reading. We first examined the temporal trend in the mean length of individuals returning for the first time to investigate a presumed decrease in body length. We then tested four non-exclusive hypotheses: whether such a decrease was due to i) long-term effects of smaller length at seaward migration in smolts, ii) fewer years spent at sea and a younger age at first return, iii) a shorter growing season at sea and an advanced date of return, and vi) a decrease in the intrinsic growth rate at sea and a smaller length-at-age. The results provide insights into the ecology of sea trout in southern Europe and illustrate ongoing changes in life history strategies.

## Materials and methods

### Study site

The Bresle River, located in northern France, drains a 742 km² catchment into the English Channel at Le Tréport harbour (Figure 1). Mean annual discharge is 7.3 m^3^ s^-1^, and salmonids can access *c*. 2/3 of its total wetted length (137 km). Human activities strongly impact the Bresle estuary, and the Le Tréport harbour is the first obstacle to migration from the sea; however, the harbour is equipped with a fishway, which provides access to the main river (Euzenat et al., 2007). The Bresle River hosts a wild population of sea trout that has a life cycle of 1, 2, or 3 years in freshwater (hereafter, FW1, FW2, and FW3, respectively) and 0, 1, or 2 years at sea until first return (hereafter, SW0, SW1, and SW2, respectively).

**Figure 1.**
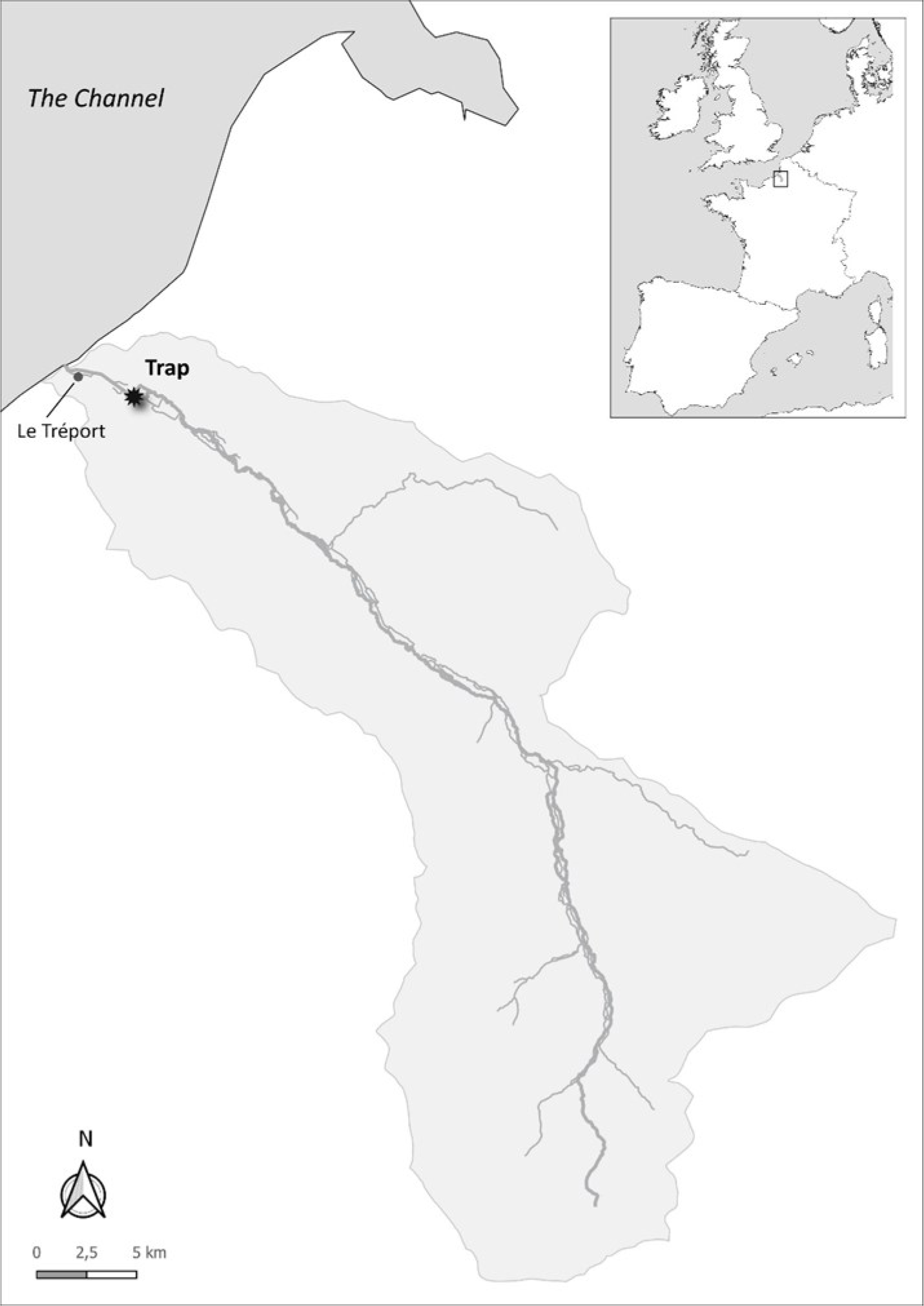
Geographical location of the Bresle River, France, and its catchment

### Monitoring protocol

Since 1981, an intensive sea trout trapping protocol has been conducted throughout the upstream migration season (March-December) each year at Eu, in the lower section of the Bresle River (Josset et al., 2023). It follows the ethical review from the committee C2EA-07, from the National Committee for Ethical Reflection on Animal Experimentation (permit number: APAFIS#26834-2020080615584492 v3). The trap is located on an obstacle 3 km upstream from the estuary and is checked twice a day for the presence of fish. Each year, a mean of 995 (SD = 511) sea trout are caught in the trap on their return from the sea. Each individual is measured to the nearest millimetre (fork length – from the tip of the snout to the fork of the tail) using a manual or electronic measuring board. Examination of secondary sexual characteristics, which are visible mainly from September-December, indicates the sex of certain individuals. However, visual sexing is imperfect and relies mainly on the presence of an elongated lower jaw (*i.e.* kype) on males. Scales are collected from a subsample of individuals (scale sampling stratified by length until 2017, all individuals sampled from 2018 onward) to determine age (all sampled individuals aged until 2012, 150 individuals aged per year from 2013 onward), following Baglinière et al., (2020). Data on individual biometry, age, and scale samples are stored in the French COLISA archive (COLlection of Ichtyological SAmples, (Marchand et al., 2019)), https://colisa.fr/) (in French).

### Data selection

For this analysis, we used data collected over a 39-year period (1984-2022). Trapping was disrupted in 2001, which reduced the efficiency and the period of operation, only SW1 individuals were captured this year. We focused on individuals that returned to the river after their first marine sojourn (hereafter, “first returns”; 75.3% of available samples). This aimed at reducing heterogeneity between individuals, as biological processes that influenced individuals returning for the first time may have differed from those that influenced individuals that had already returned to the river at least once. The returning status was defined based on the examination of scales, which was available on a subset of 15,730 individuals out of the 41,412 individuals captured at the trap. First return fish were defined as individuals with no spawning mark on their scales. Individuals with low relative weight (Wr < 90) (Blackwell et al., 2000) captured before April were considered late-running post-spawners migrating to the sea and were thus excluded from the analysis.

### Data resampling

The subsample of fish which had their scales sampled, and age assessed, was unbalanced between length classes, with individuals in the more abundant length classes being generally under-sampled. This bias skewed estimates of the mean length-at-age. To correct this bias, we randomly resampled within the subsample of fish which had their scales sampled, a subset of individuals with an even effort among length classes. We first calculated the proportion of sampled individuals for each 50 mm length classes for a given year, determined the minimum proportion of sampling for each year, and then used this minimum proportion as a target for resampling individuals in each length classes for that given year. This resampling procedure was repeated 1000 times to ensure robustness and stability of the results. The following selection of first returns, yielded datasets of varying lengths, with a mean sample size of 6060 individuals (SD = 23).

### Statistical analyses

#### Long-term trend in the length of returning sea trout

To describe the long-term temporal trend in the length of first-time returning sea trout, we modelled the mean yearly length of first returns as a linear function of time and assessed the significance and direction of the trend. This was done using the “lm()” function of the “stats” package of R software (version 4.0.3) (R Core Team, 2020), using the year of return as a continuous variable.

#### Drivers of change in length-at-first return

We then investigated the influence of key ecological variables on the length of first-time returning sea trout using Gaussian generalized linear models (McCullagh and Nelder, 1989) with a log link. The log-link function allows for multiplicative rather than additive effects, and the former are generally considered more appropriate for modelling ecological relations. The Gaussian distribution was selected because it is usually used when a linear relation between dependent and independent variables is expected, assuming a normal distribution of the residuals. Model selection followed stepwise inclusion of variables and relevant two-way interactions depending on the hypothesis selected.

Development of dimorphic sexual characteristics before reproduction, especially a kype at the end of the lower jaw (Witten and Hall, 2001), could bias length depending on the timing of return. This would be especially true in late-running males, although some females also have a small kype. To address this bias, we used the apparent sex recorded in the phenotypic data in the model as a corrective variable, with two levels: presence or absence of a kype.

Knowing that the longer the marine sojourn, the larger the trout is upon return (Thorstad et al., 2016), we included the “sea age” (*i.e.* the number of years spent at sea) as a categorical variable to explain length-at-first return. Similarly, we hypothesised that the number of years spent in the river positively influenced the length-at-first return, although less than the number of years spent at sea, due to substantial differences in growth potential in fresh and sea-water (Gross et al., 1988; Nevoux et al., 2019). We thus used the “river age” (*i.e.* the number of years spent in freshwater before migrating to the sea) as another categorical variable in the model. We also considered an interaction between river age and sea age to capture effects of latent intrinsic differences among individuals depending on their life history (*e.g.* growth potential, metabolic rate, energy-use efficiency). Temporal variability in the effect of age on length was also tested by considering interactions between the year and river and sea ages.

The timing of migration can also influence the length of returning trout. Assuming that the timing of smolt migration to the sea remained unchanged (De Eyto et al., 2022), an early return in freshwater would mean shorter marine sojourn, reduced growth opportunities and potentially shorter bodies. Advanced migration timing can be driven by environmental conditions experienced by all individuals of a given sea-age class, as indicated by the mean day of year of return of each sea-age class and year (ctrAvgDOY, centred on the inter-annual mean). Within a given sea-age class, differences in the timing of migration among individuals can also influence body length. This individual level was captured by the variable deltaDOY, which was calculated as an individual’s return day minus the mean day of return of the corresponding sea-age class and year. We considered two-way interactions to test whether effects of phenology variables varied over time or among sea ages. The increase in length for each additional day at sea may have varied during the study period, and timing of return may have more influence on body length of young individuals returning after a short sojourn, due to their faster growth (Davaine et al., 1997).

Once the migration strategy and timing were considered, any remaining long-term trends in the length of returning sea trout would indicate a gradual change in the intrinsic growth rate at sea. To describe residual temporal variability in the length-at-first return, we used the year of return to the river (Year) as a continuous variable centred on 1980, which grouped sea trout that had experienced the same environmental conditions during their last months at sea.

#### Implementing and selecting the model

Collinearity among variables was tested using Pearson correlation coefficients, ensuring that the correlation coefficient between predictor variables |r| remained inferior or equal to 0.7 following Dormann et al., (2013). A bootstrap procedure of the “boot” package (Canty and Ripley, 2021; Davison and Hinkley, 1997) was used to ensure stability of the results by reducing effects of sampling variability. One thousand datasets were generated using the bootstrap procedure, and candidate models were fitted to each dataset. Model selection aimed to minimise Akaike’s information criterion (Akaike, 1974), following guidelines of Burnham et al. (2002). We identified the best model for each dataset and compared the models’ selection frequencies as a measure of support. Mean estimates and 95% confidence intervals of the best models’ effects and interactions were then calculated from the 1000 iterations of the bootstrap procedure and presented relatively to a reference SW1-FW1 sea trout with a kype and returned on the average day of return of its sea age class. In addition, to assess temporal changes in the age structure, yearly age-class percentages were recorded for each of the 1000 iterations. Similarly, slopes of a linear regression between the day of year and the year of return by sea age were extracted from each iterations to assess variations in the timing of return.

## Results

### Long-term decrease in the length of returning sea trout

The temporal regression on the body length of first returns from 1984-2022 revealed that first-time returning sea trout in the Bresle River lost on average 1.73 mm (SD = 0.08) per year, regardless of their age, which represented a loss of 67.47 mm (SD = 3.12) (*i.e. c.* 12.3% of the mean body length from 1984-1988) over the 39 years studied (Figure 2).

**Figure 2.**
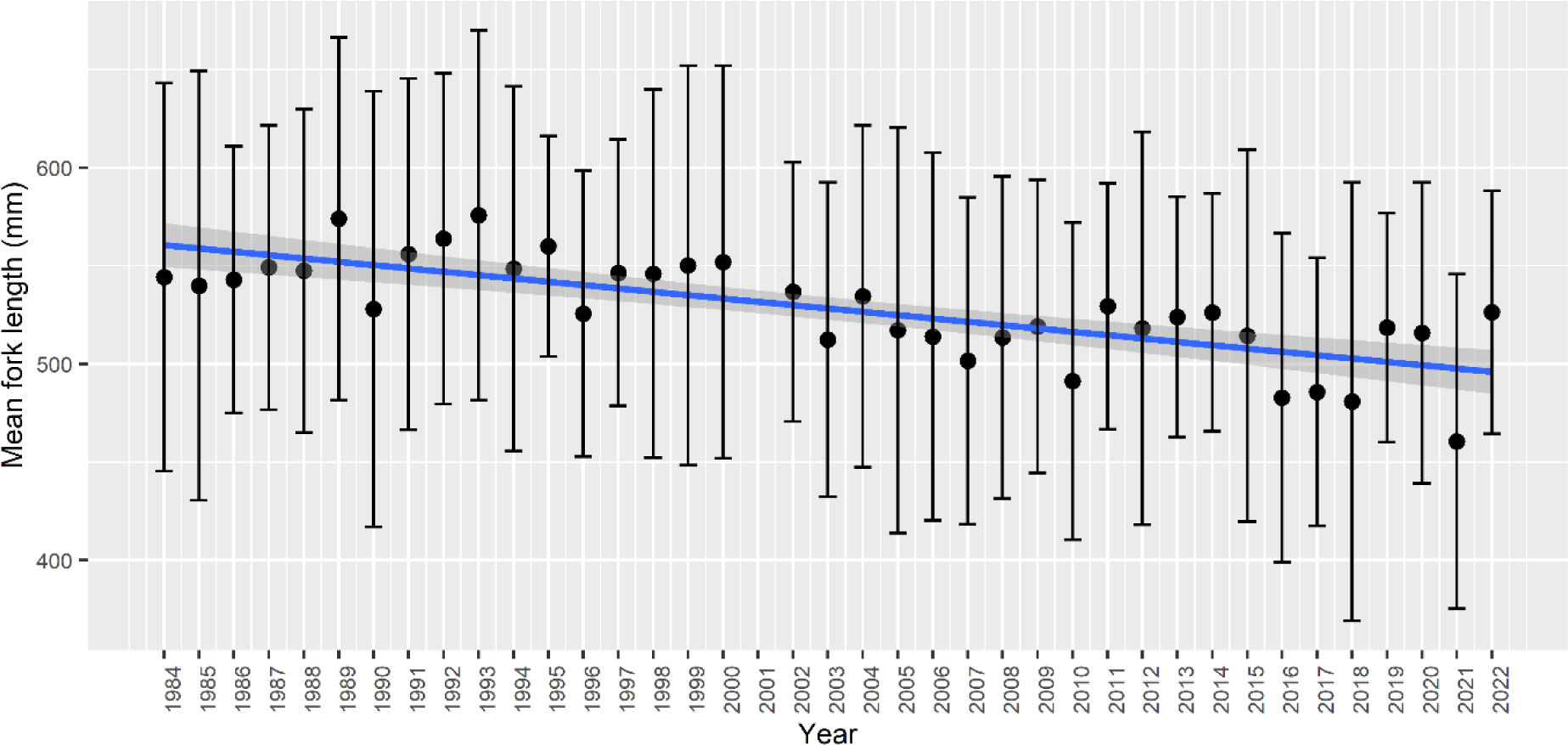
Long-term temporal trend in the mean fork-length of first-time returning sea trout captured in the Bresle River from 1984-2022. Error bars indicate 1 standard deviation. The shaded zone indicates the 95% confidence interval. Year 2001 is not presented due to floods that disrupted the trapping, which lead to low efficiency and the capture of only SW1 individuals.

### Model selection

Of the 1000 iterations of the bootstrap procedure and model selection, only two models (no. 22 and 20) were selected for having the lowest AIC (selected 77.3% and 22.7% of the time, respectively) (Table 1). The two models were similar, including all major variables, and thus had similar AIC and predictive power. They differed only in that model 22 included temporal interactions with the two phenology variables (*i.e.* ctrAvgDOY×Year and deltaDOY×Year).

**Table 1.**
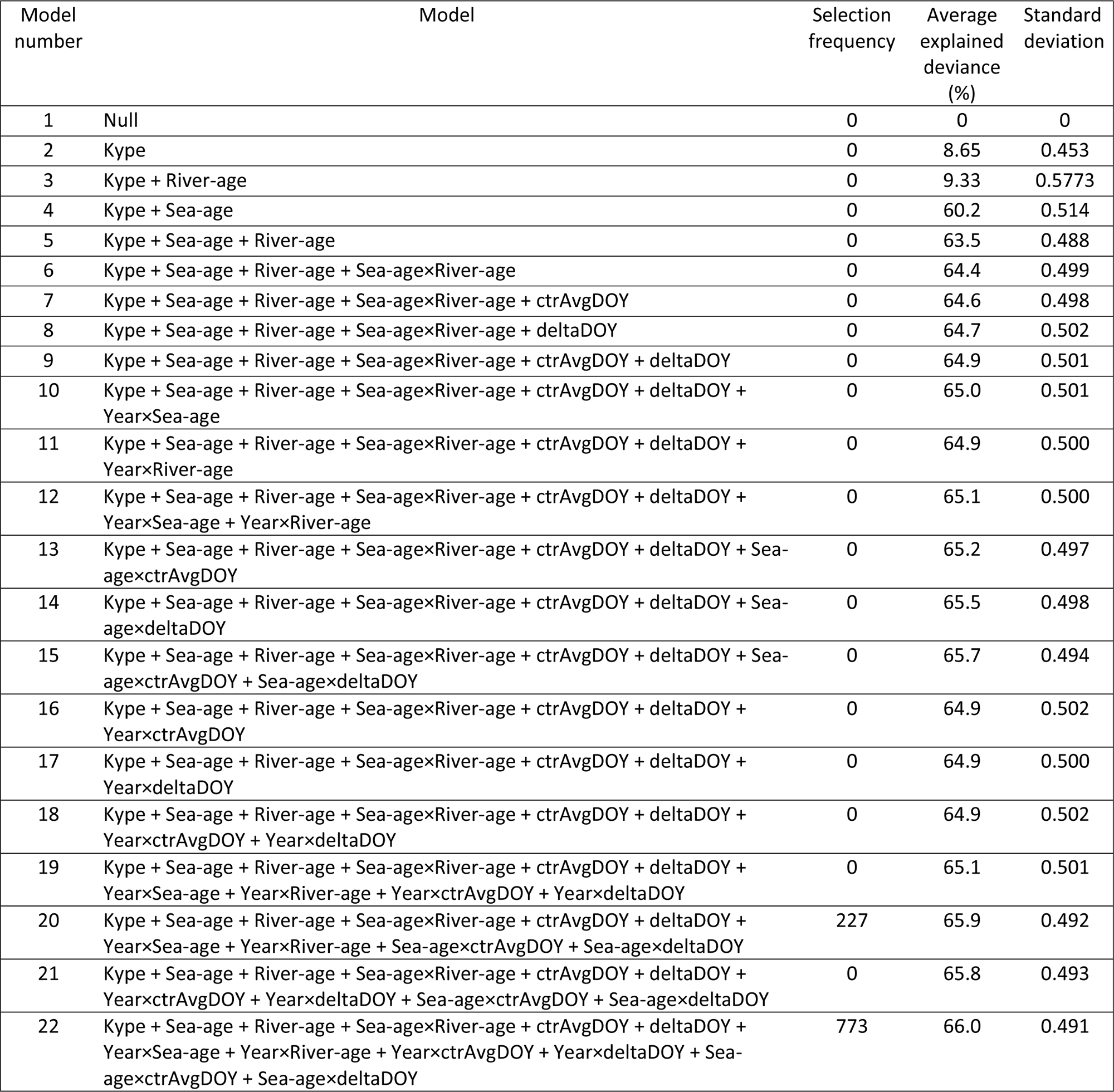
The models tested to explain length-at-first return and their frequencies of selection (out of 1000 iterations) based on Akaike’s information criterion. FL, fork length; Kype, presence/absence of a kype; Sea-age, number of years spent at sea; River-age, number of years spent in the river; ctrAvgDOY, mean day of year of return to the river by year and sea age (centred by sea age); deltaDOY, individual difference in the mean day of year of return; Year, year of return to the river.

Although model selection based on the AIC balances likelihood and parsimony, two of the most complicated models were selected, which indicates overall complexity of the mechanisms that drive length-at-first return. Presence of a kype positively influenced the length at first return and explained up to 8.65% (SD = 0.453) of explained deviance alone. The mean deviance calculated from the 1000 best models explained up to 66.00% (SD = 0.49) of the variability in the length-at-first return. Examination of the residuals showed slight deviation from the hypothesis of normality. As could be expected in a large dataset, wide tail distributions lead to the rejection of the normality hypothesis despite a quasi-normal distribution.

### Decrease in the age of returning sea trout is a major driver of decrease in length-at-first return

We found that sea age was the most influential variable on length-at-first return, as it lead to the highest gain in explained deviance (Table 1). If we compare the effect of variables, averaged over the 1000 best-models, we show that the more growth seasons an individual spent at sea, the greater was its length-at-first return (Figure 3). Returning after 2 years at sea lead to a 30.08% increase (95% CI [30.03; 30.1]) in length-at-first return compared to the reference fish, while returning as SW0 lead to a 37.53% decrease (95% CI [-37.59; -37.48]). This result supports the hypothesis that a shift in the age structure towards younger individuals could drive the decrease in length observed among first returns. Indeed, the mean sea age structure of first returns changed greatly during the study period (Figure 4). In particular, 2SW individuals (293 individuals in average across the 1000 datasets, SD = 67) gradually disappeared and were not represented in 3 of the last 10 years of the study period.

**Figure 3.**
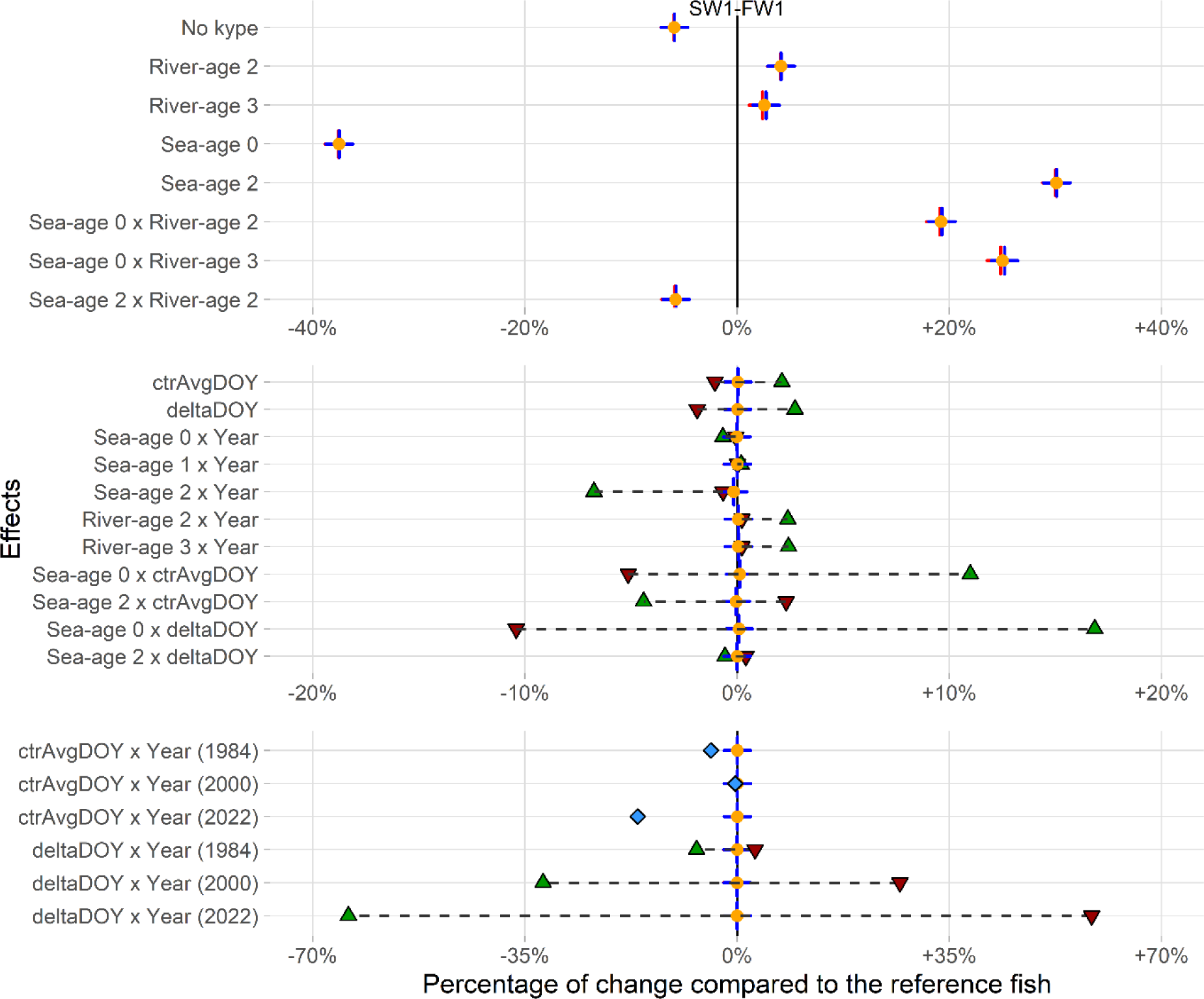
Mean estimated effects on the length at return in percentage of change, averaged over the 1000 best models and compared with a reference SW1-FW1 sea trout, with a kype, for ctrAvgDOY and deltaDOY = 0 (black vertical line). Top panel: effects of qualitative predictors and their interactions, middle panel: effects of quantitative predictors and interactions between quantitative and qualitative predictors, bottom panel: discretised effects of interactions between quantitative predictors. Orange dot, red and blue crosses indicate respectively mean estimated effect, lower and upper 95% Confidence Intervals (See Table 1 for the definition of variables.). Downward red triangle and upward green triangle indicate the mean estimated effects when the quantitative effect is at its minimum and maximum values respectively for the reference sea age class (SW1); dashed black lines illustrate the range of possible values. Blue diamond indicate the mean predicted effect for the discretized ctrAvgDOY:Year interaction for the reference sea age class, i.e. 1 value per year per sea age class. Effects are presented in the ‘response’ scale, i.e. -10% indicate that an individual with the specific modality would be estimated to return shorter by 10% compared to the reference fish.

**Figure 4.**
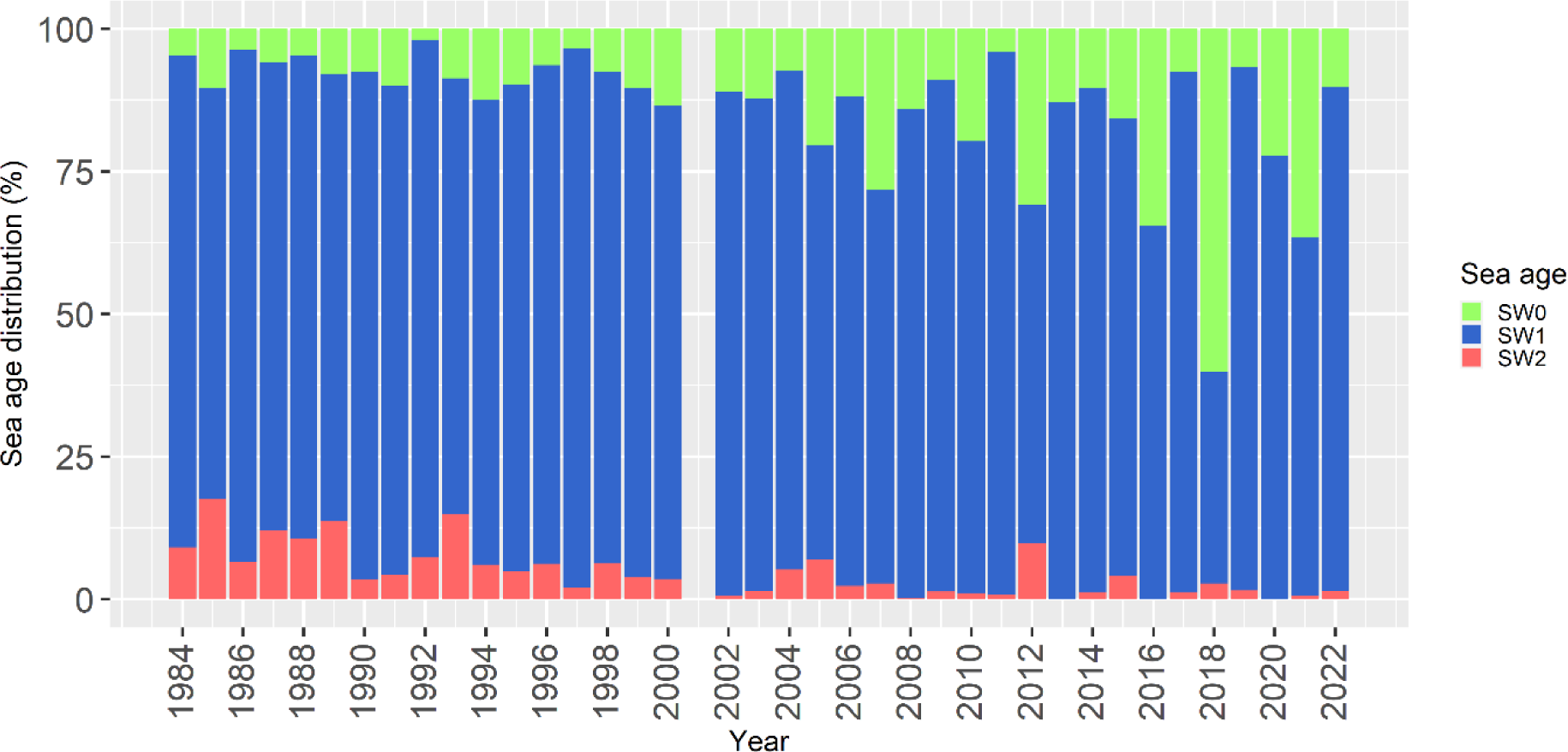
Temporal variability in the age structure of first-time returning sea trout captured in the Bresle. Average proportions derived from the 1000 datasets. Year 2001 is not presented due to floods that disrupted the trapping, which lead to low efficiency and the capture of only SW1 individuals.

At the same time, the percentage of SW0 individuals increased in recent years to up to 60% (in 2018) of all first returning sea trout. The percentage of SW1 individuals in the population remained predominant and stable over time (mean = 81.8%; SD = 11.2%). This pattern indicates a decrease in the mean sea age at first return of the Bresle sea trout population: 1.06 years (SD = 0.4) from 1984-1988, but 0.80 years (SD = 0.4) from 2018-2022.

### Earlier life history contributes to the length of returning sea trout

River age had a long-term effect on the length-at-first return: for a fish that spent one year at sea, individuals that smoltified after only 1 year in the river (the reference SW1-FW1 individual) were shorter on their return than those that smoltified after 2 or 3 years, which were predicted to be respectively 4.13% (95% CI [4.11; 4.15]) and 2.55% (95% CI [2.38; 2.72]) larger (Figure 3). The younger the sea-age class, *i.e.* the shorter the time spent at sea, the more pronounced this effect was, as indicated by the interaction between river age and sea age. For instance, a SW0 individual smoltifying at age 2 or 3, was predicted to be respectively 19.22% (95% CI [19.13; 19.31]) and 25.02% (95% CI [24.83; 25.22]) larger than the reference SW1-FW1 fish (Figure 3). River age was included in all of the best models, which highlights its high explanatory value.

### Length of returning sea trout is influenced by the migration timing

The two variables for migration timing (ctrAvgDOY and deltaDOY) explained relatively small amount of the variability in the length-at-first return. As displayed on Figure 3 by minimum (downward red triangles) and maximum (upward green triangles) values, their main effects ranged respectively between [-1.05%; 2.1%] and [-1.88%; 2.73%]. Effects of return timing depended on the sea age, as indicated by the inclusion of the two-way interactions in the model. The effect of additional days at sea, measured at both inter-annual (ctrAvgDOY) and intraannual (deltaDOY) scales, seemed to benefit SW0 individuals more than SW1 individuals, with a maximum of +16.9% effect for the Sea-age 0 x deltaDOY interaction. In contrast, this effect was negative for SW2 individuals, as late-returning cohorts in this age class tended to be smaller (-4.4% at most) than early-returning ones. The inclusion of the temporal interaction indicated a potential change in effects of return timing over time, although support for it was weaker, as 22.7% of the best models did not include it. The range of theoretical effect for these interactions was wide, with for instance the deltaDOY x Year interaction for 2022 ranging between [-64.1%; +58.5%], although extreme values were actually rare and opposing effects (deltaDOY main effect, Year in interaction with other variables) must be taken into account.

Mean slopes of temporal linear regressions of the day of return indicated that the capture date of first returns advanced significantly (p < 0.001), with SW0, SW1, and SW2 being captured a mean of 1.38 (SD = 0.12), 1.21 (SD = 0.08), and 0.49 (SD = 0.26) days earlier each year, respectively. Over the 39-year period, the date of return thus advanced by 53, 47, and 19 days for SW0, SW1, and SW2 individuals, respectively.

### Weak and contrasting evidence of changes in growth

Interactions between year and both river age and sea age were included in the best models, which supports the hypothesis of temporal changes in the length-at-first return as a function of life history strategy. The effect of the year on length-at-first return remained small and relatively stable for FW1, but slightly increased over time for FW2 and FW3, ranging respectively between [+0.22%; +2.3%] and [+0.22%; +2.4%] (Figure 5), both of which had similar variations (Figure 5). Furthermore, length-at-sea-age over time differed, being stable or slightly positive for SW0 and SW1 individuals, but decreasing for SW2 individuals, accounting for a decrease of -6.7% relative to the reference individual in 2022. Overall, except for SW2 individuals, trends remained weak, with only slight changes in the length-at-age for SW0 and SW1 individuals.

**Figure 5.**
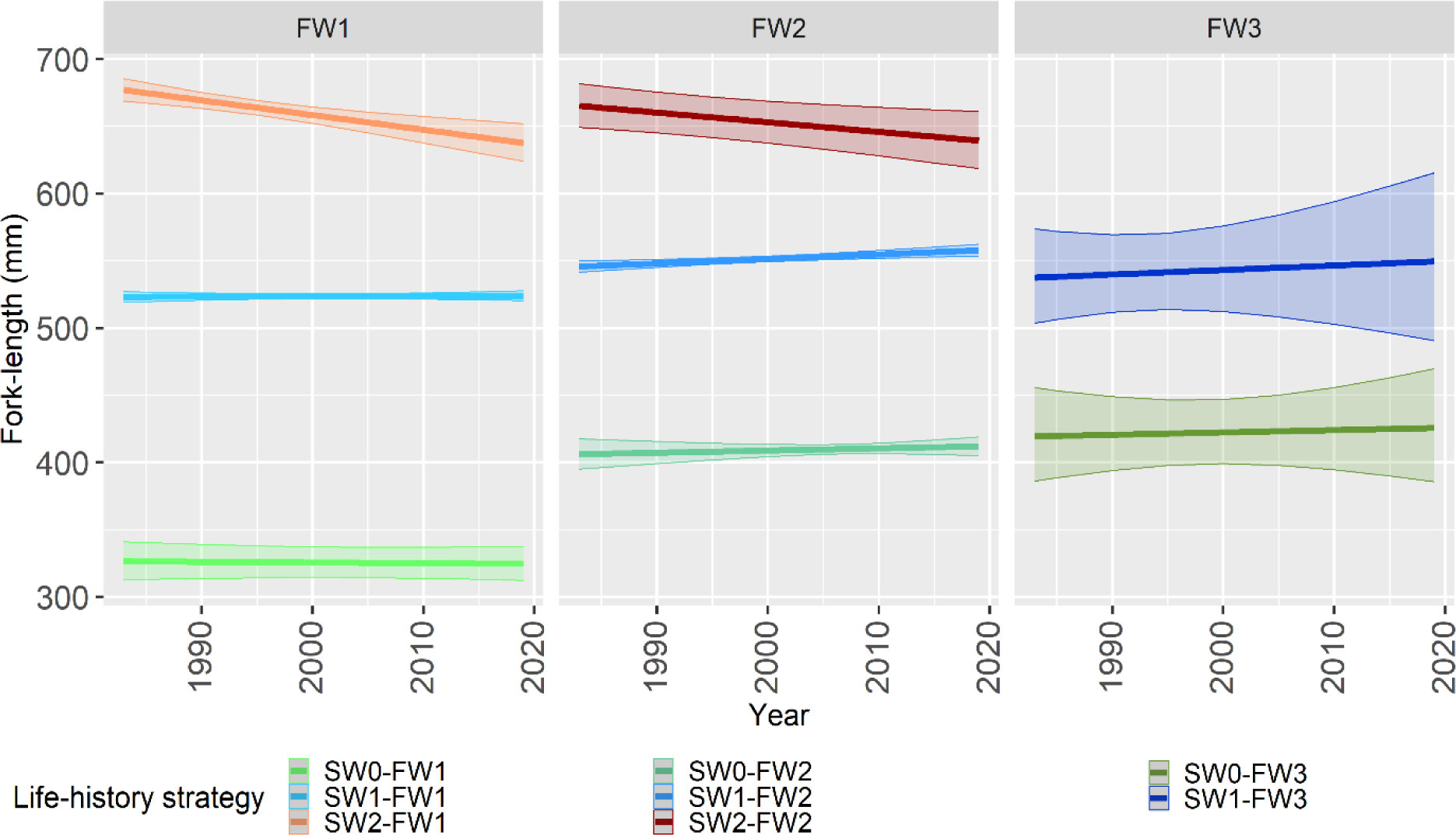
Estimated changes in the length of sea trout at first returns over time as a function of life history strategy. Mean fitted values in the ‘response’ scale derived from predictions on each best models. Shaded areas indicate 95% confidence interval. FW: number of years spent in river, SW: number of years spent at sea. For readability, plots are presented by river age. Wide confidence intervals for FW3 can be explained by the relatively low number of individuals (20 individuals on average over the 1000 datasets; SD = 9).

## Discussion

Results indicate that individuals returning to the Bresle River for the first time became progressively smaller over the 39 years of the study period. This decline mainly resulted from change in the age structure and contrasted variations in length-at-age. In the study population, an increasing percentage of sea trout returned for the first time at age SW0 and the return migration timing advanced. This decrease in duration of the marine sojourn can explain the decrease in body length, while length-at-age remained constant or slightly increased over time for SW0 and SW1 individuals. At the same time, the percentage and length-at-age of SW2 individuals decreased.

### Strengths and limitations of long-term datasets

Sampling biases are inevitable during long-term monitoring of wild populations. It is often challenging to maintain constant sampling effort over long periods, whether due to changes in the sampling protocol, a lack of resources, modifications of the observation device or objectives, or varying environmental conditions. In this study, the need to control for sampling biases required an analytical rebalancing of the dataset. Using the bootstrap procedure enabled multiple iterations of subsampling and modelling, which increased the stability and robustness of the results. One downside of such large datasets is the risk of rejection of the assumption of normality in the residuals, as required by GLMs. Non-Gaussian distributions, such as the Gamma distribution, may reduce this risk. However, this option did not improve the fit of the residuals in the present case. McCullagh and Nelder (1989) reported that a rejection of this assumption was not of major importance in the case of “large datasets”, where minor deviations from the normality will likely lead to rejection of the assumption. Deviations in the residuals suggests either errors in the determination of age by experts, or high heterogeneity in growth performance between individuals that is not captured by the model. Sea trout being famous for the extreme plasticity in its life history traits and physiological performance (Nevoux et al., 2019; Thorstad et al., 2016), the latter is likely to be an important factor here. Despite these difficulties, long-term datasets are crucial for understanding highly complex ecological and evolutionary processes (Eliott and Eliott, 2006; Hughes et al., 2017).

### Decrease in mean population length does not necessarily imply a decrease in length-at-age

With a decrease of *c.* 12.3% in first-time returning sea trout body length over the past four decades, the results add to the growing corpus of literature that describes a decrease in the length of salmonid species around the world (Bal et al., 2017; Jeffrey et al., 2017; Lewis et al., 2015; Morita and Fukuwaka, 2007; Ohlberger et al., 2018; Oke et al., 2020) or in other fish families (Boëns et al., 2021; Neuheimer and Taggart, 2010). The extent of the changes observed in the Bresle sea trout population are slightly higher than results of Oke et al. (2020) for four species of Pacific salmon (Chinook, *Oncorhynchus tshawytscha*: -8.0%, coho, *Oncorhynchus kisutch*: -3.3%, chum, *Oncorhynchus keta*: -2.4%, and sockeye, *Oncorhynchus nerka*: -2.1%), although the underlying mechanism may differ. While *Onchorhynchus* species may suffer from a reduction in individual growth rate at sea, interestingly in sea trout we highlighted that individual length-at-age changed little overall in most age-classes. A decrease in the age at first return appears to be the main reason of the decrease in the length-at-first return observed for sea trout in the Bresle River. Structural changes in the age at return of sea trout have previously been reported by Milner et al., (2017), who found a change in the proportion of low-weight fish in rod catches of four Welsh rivers between 1977 and 2007, suggesting an increased proportion of SW0.

It was surprising to detect a clear negative temporal trend in the length-at-age only for SW2 first returns, while those of SW0 and SW1 remained relatively stable. Marine trophic webs are under tremendous pressure in the context of climate change. The general consensus is that ocean productivity has decreased in the past few decades and that the composition of its ecosystems has changed greatly (Behrenfeld et al., 2006; Gohin et al., 2019; Hoegh-Guldberg and Bruno, 2010; Möllmann and Diekmann, 2012). One would thus expect to observe a decrease in growth in all age classes, and in younger age classes in particular, as observed in Atlantic salmon (*Salmo salar*) populations returning to nearby rivers in France and Southern England (Tréhin et al., 2021). However, our results suggest that SW0 and SW1 could sustain stable growth rate over the 39-year study period. In fact, Davidson et al., (2006) even reported increased post-smolt growth for these two age-class in the sea trout population of the River Dee. This differing response among sea-age classes could indicate local changes in the conditions that sea trout encountered at sea, rather than a broad decrease in growth opportunities. During migration at sea, salmonids gain access to different feeding grounds and sets of prey (Knutsen et al., 2004, 2001; Rikardsen et al., 2006, 2007; Thorstad et al., 2016). Consequently, changes in food availability and quality on the feeding grounds used by older and larger sea trout may explain their lower growth. Additionally, local variations in sea temperatures could also influence sea trout return size, as reported before by Milner et al., (2017) in the Irish sea.

### Potential drivers of a change in sea trout age structure, and demographic consequences

A decrease in marine survival can select for a reduction in the duration of the sea sojourn and early maturation to maximise fitness (Archer et al., 2019; Thorstad et al., 2016). Parasitism by sea lice (*Lepeophtheirus salmonis*, Krøyer, 1837) has been demonstrated to be a major factor of reduced survival in wild salmonids (Bjørn et al., 2001; Thorstad et al., 2015) and is considered to as a major threat for many sea trout populations in Norway (Fiske et al., 2024) and Ireland. Sea lice may have been responsible for the collapse of the sea trout population in the Burrishoole River (Gargan et al., 2006) and for profound changes in life history traits of that in the Eriff River, Ireland (Gargan et al., 2016). In the Bresle River, most returning sea trout show evidence of sea-lice parasitism, with the youngest and smallest individuals usually having the highest parasite loads. Premature returns, in which non-mature individuals with heavy sea-lice loads return to freshwater for delicing, have been well documented in the literature (Birkeland, 1996; Birkeland and Jakobsen, 1997; Serra-Llinares et al., 2020). Thus, parasitism and the related increase in the returns of young non-mature individuals could provide an explanation for the decrease in the average sea age of first returns, as observed in this study.

The lack of data about the reproductive status of first-time returning individuals limits our understanding of the role of SW0 in the population dynamics. The return status of the individuals included in this study was determined from the analysis of patterns on scale reading. However, it provides an imperfect record of previous reproduction history (Baglinière et al., 2020). Being able to separate out the mature SW0 returns from the immature SW0 individuals (Birkeland, 1996) would help better predict the impact of changes in sea-age composition on population dynamics. In the nearby Calonne River, while most SW0 individuals were mature in the 1980s (Maisse et al., 1991), recent changes in environmental conditions, and sea-lice prevalence in particular, may have profoundly altered maturation and return strategy.

Furthermore, age at maturation has been demonstrated to be a partially heritable life history trait in sea trout (Ferguson et al., 2019; Reed et al., 2019). Thus, natural selection and change in allele frequency could potentially explain the progressive shift in sea-age structure toward shorter sea sojourn observed in this study. Length-specific selective mortality that targeted mainly larger individuals (Reznick et al., 1990) could as well explain the progressive disappearance of SW2 individuals in the Bresle population. Overfishing has been widely demonstrated to influence many characteristics of fish populations, such as length at reproduction (Neuheimer and Taggart, 2010; Ojuok et al., 2007) and a direct selection on length is not necessarily required to result in changes of age and size at first return (Hard et al., 2008; Kozlowski, 2006). Intensive net fishing, which is typically size-selective, occurred at sea near the Bresle estuary, especially during the first half of the study period. With up to 37% of the population being caught each year either by professional or leisure net fishing at sea or rod-and-line fishing in the river (Fagard and Beaulaton, 2018), it seems plausible that it could contribute to a reduced age at first return, as observed here. Besides, the resurgence of marine apex predators has been associated with a decrease in the mean length of Chinook salmon (Ohlberger et al., 2019). However, although the Bresle River is located near the Bay of Somme, which hosts the largest breeding colony of harbour seals (*Phoca vitulina* L.) in the English Channel, analysis of 86 scat samples from this colony revealed no evidence of salmonids in the seals’ diet (Spitz et al., 2015), suggesting that salmonids do not contribute significantly to it. Furthermore, most seal predation on salmonids seems to be opportunistic, with no indication of length-dependent selection for larger fish (Suuronen and Lehtonen, 2012; Thomas et al., 2017). Therefore, it seems unlikely that seal predation would have driven the population-wide changes in sea trout age structure observed here.

Changes such as the progressive loss of a sea-age class, as observed for SW2 individuals in this study, can strongly influence a population’s resilience (Erkinaro et al., 2019; Stewart, 2011). The loss of older age classes can also have an especially detrimental effect due to the loss of “big, old, fat, fecund, female fish” (BOFFFFs). These older female fish contribute greatly to reproduction as they produce more and often larger eggs than younger females do (Hixon et al., 2014; Ohlberger et al., 2020). This reproductive hyperallometry has been demonstrated to be widespread both in fishes and other taxa with indeterminate growth, such as mollusc or crustaceans, and suggests that the reproductive role of these large and older females has been widely underestimated (Barneche et al., 2018; Marshall and White, 2019). Therefore, management actions such as specific fishing regulations to protect these older fish could be implemented in the Bresle River.

### The importance of considering the entire life history

When sea age was taken into account, river age also had a positive effect on length-at-first return, which suggests that length at migration had long-term consequences until the first return. Growth potential at sea is much higher than it is in freshwater, accordingly the latter often seems to be viewed as a mere transitional habitat between hatching and smoltification. Nevertheless, the hypothesis of a mechanistic length threshold driving maturation decision (Hutchings and Jones, 1998; Tréhin et al., 2021) predicts that larger smolts will reach the critical maturation length sooner and thus return after fewer years at sea (Tattam et al., 2015). Therefore, this carry-over effect of a trout’s freshwater life on its length-at-first-return hold important implications for an individuals’ future life history. It also suggests the importance for managers not to neglect actions aimed at protecting juveniles. In particular, these action should focus on the preservation of phenotypic composition (age and size structure)(Russell et al., 2012) in a context of climate change. These actions could especially include managing and protecting water resources (Waco and Taylor, 2010), restoring habitat connectivity (Forget et al., 2018), and creating thermal refuges (Blann et al., 2002). Our results also suggests that the effect of river age on length at return is most visible in SW0 and tends to decrease as time at sea increases. This is likely a consequence of higher growth opportunities at sea (L’Abee-Lund et al., 1989) that would flatten length differences acquired in freshwater over time.

### Effect of migration timing on length-at-first return

The effect of a longer sea sojourn were positive and stronger for sea trout that returned after SW0, which is consistent with brown trout growth curves, which have steeper slopes for younger age classes before reaching a plateau (Davaine et al., 1997). Variations of the return date was important over the study period, especially in SW0 and SW1 (resp. 53 and 47 days earlier in 39 years). This observation is consistent with, but larger than, the advance of 2.6 days per decade observed by Legrand et al. (2021), in the migration timing of sea trout at 40 monitoring stations in France. Such a difference may result from trend variations across monitoring stations, as the Bresle River was the northernmost one, as well as averaging between age classes. Nevertheless, despite an earlier arrival of the order of a month, timing of return had a relatively small influence on length at first return overall. This result is concordant with a study from the River Dee, where length at return of SW0 individuals was found to be independent from the day of return (Celtic Sea Trout Project, 2016).

## Acknowledgements

Long-term datasets that support studies such as the present work would not exist without the continued work and support of successive generations of field workers, technicians, researchers, and managers, and we thank them all. We thank Fabien Quendo for the making of Figure 1. We thank PCI recommender Aleksandra Walczyńska for her comments and handling of this paper, as well as Jan Kozlowski and an anonymous reviewer for their careful reading and useful comments.

## Data and code availability

Data analysed in this study is available at: https://doi.org/10.57745/FPHLBT

Code for the analysis can be found at: https://doi.org/10.5281/zenodo.10199683

## Author information

### Authors’ contributions

**Q.J.**: conceptualization, data curation, formal analysis, investigation, methodology, validation, visualization, writing the original draft, writing the review and editing;

**L.B.**: conceptualization, funding acquisition, methodology, project administration, resources, supervision, validation, visualization, writing – original draft, writing the review and editing; **A.R.**: conceptualization, project administration, supervision, validation, visualization, writing the review and editing;

**M.N.**: conceptualization, methodology, project administration, supervision, validation, visualization, writing – original draft, writing the review and editing.

## Appendix

**Appendix 1.**
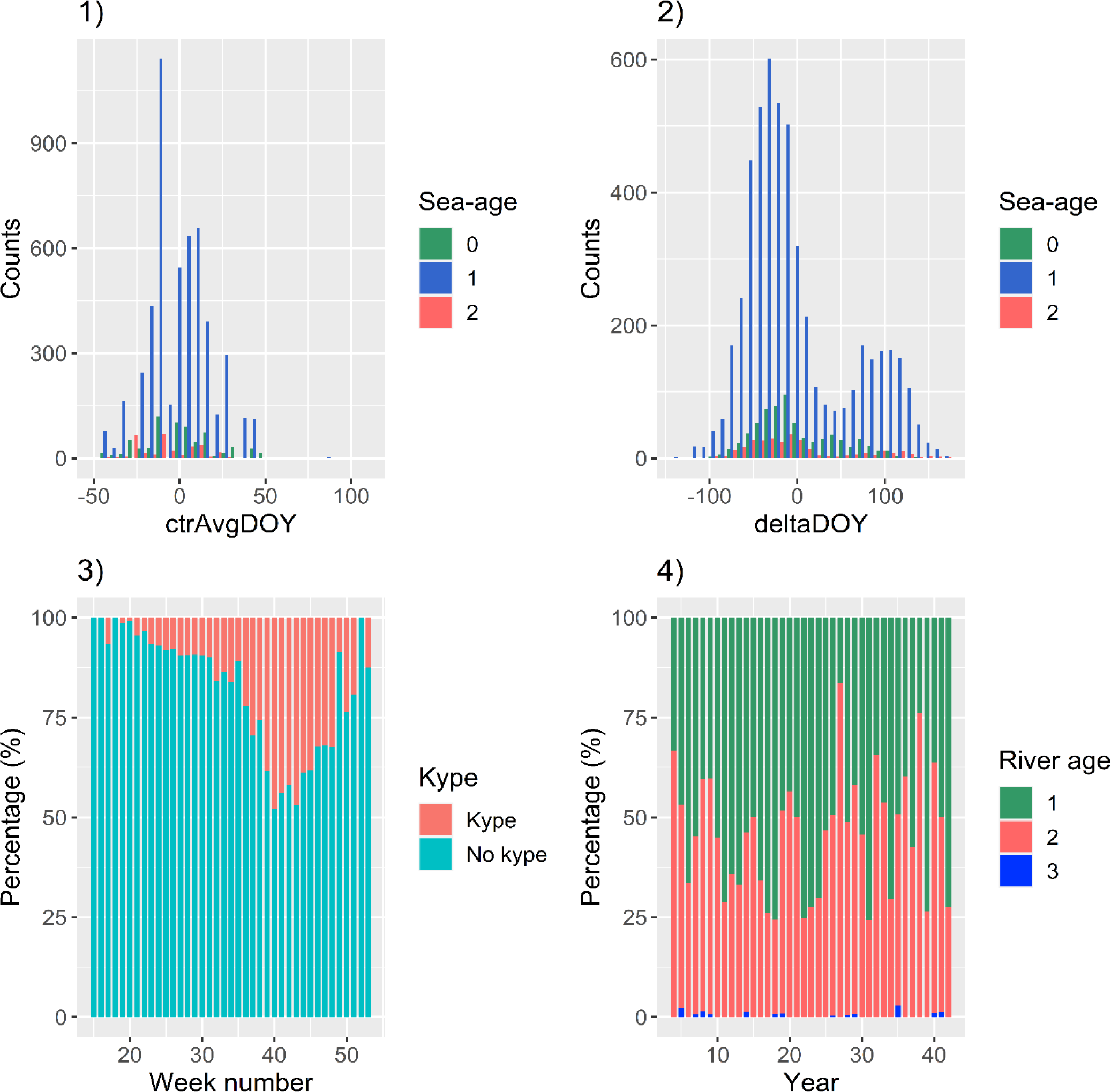
1) Histogram of ctrAvgDOY, 2) histogram of deltaDOY, 3) proportion of fish with a kype and without across weeks, 4) proportions of river ages, for one random dataset.

**Appendix 2.**
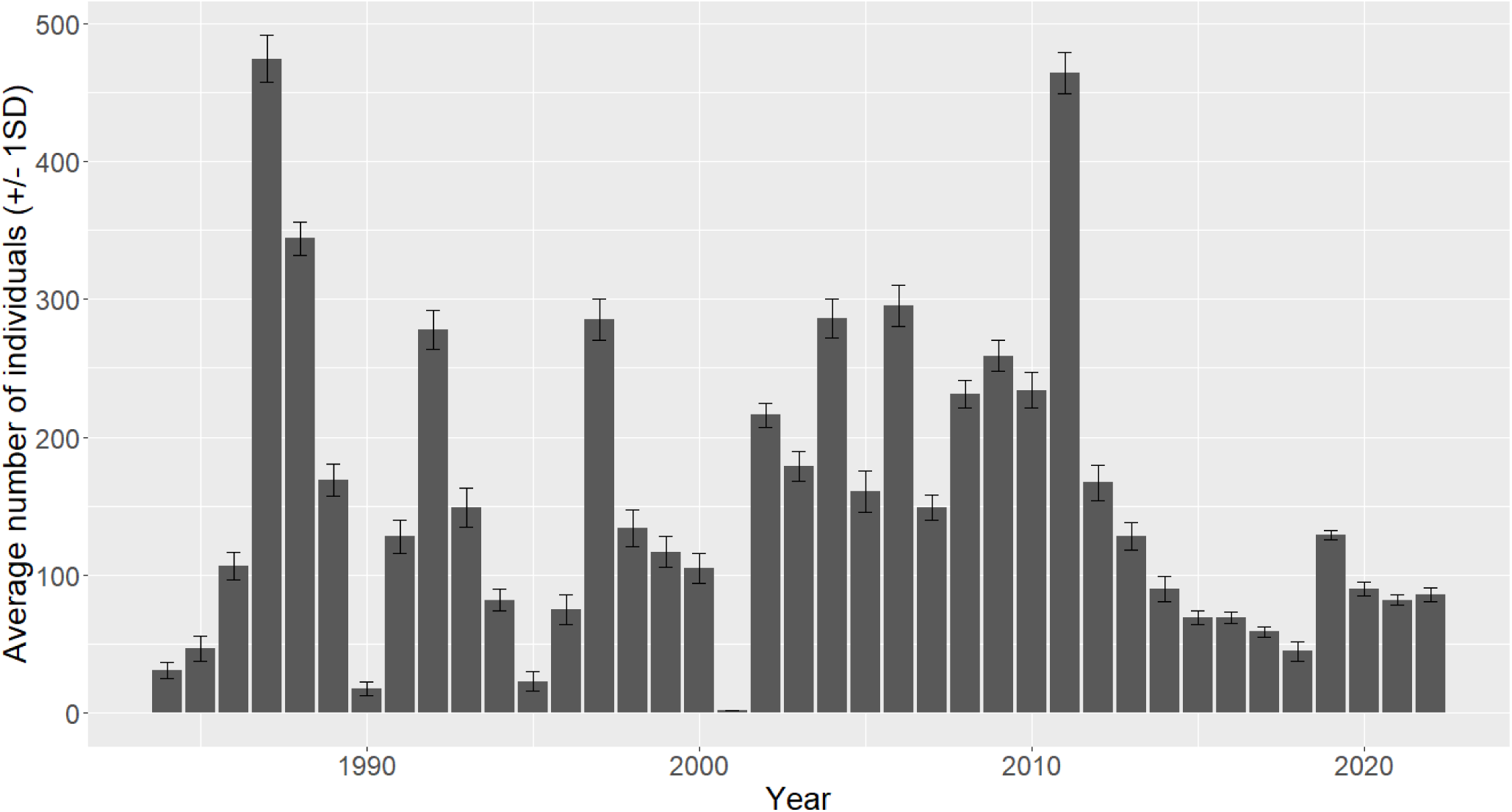
Average number of individuals per year (+/- 1SD) in the 1000 datasets.

**Appendix 3.**
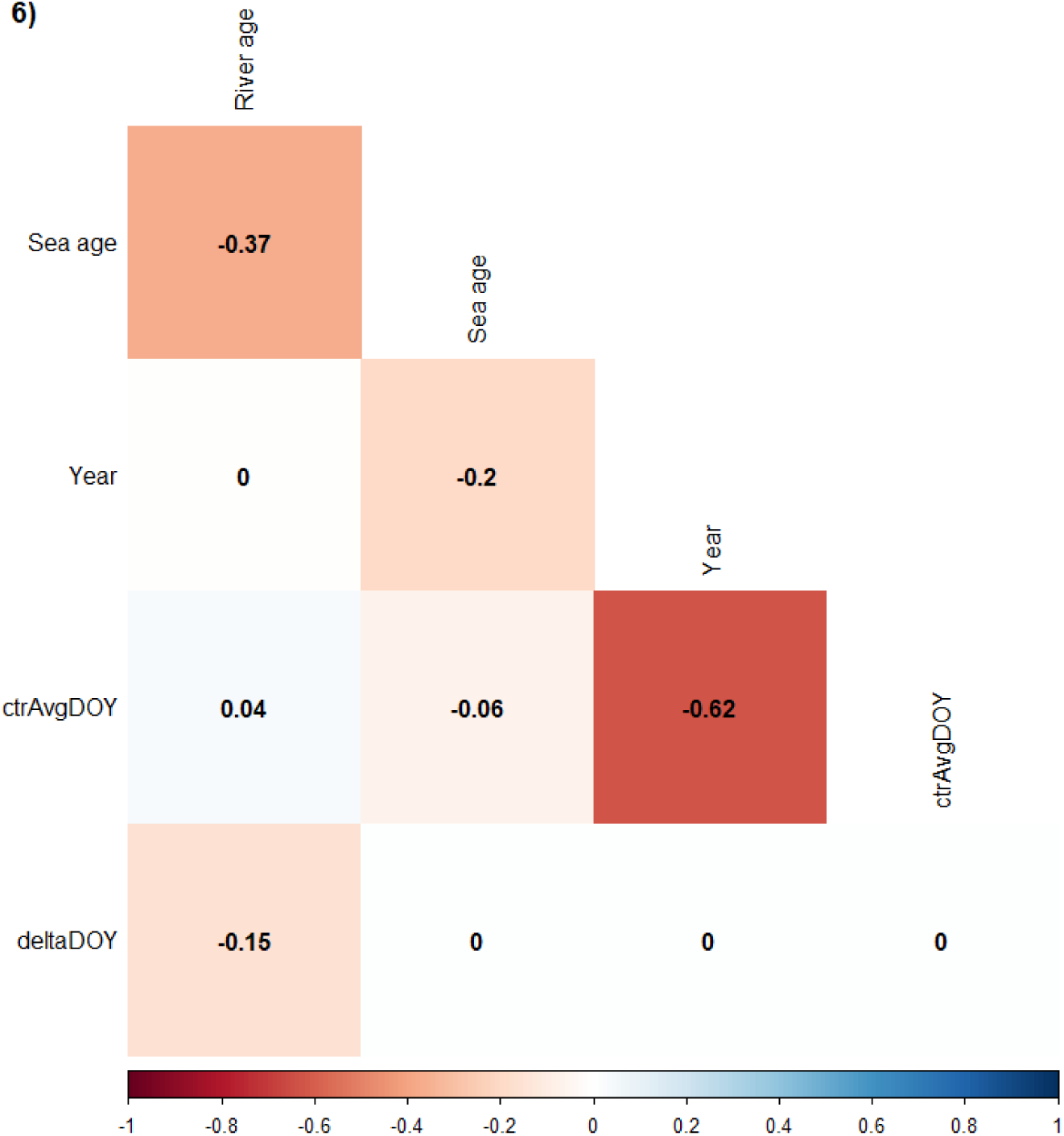
Correlation plot between explanatory variables.

## References

Akaike, H., 1974. A new look at the statistical model identification. IEEE Transactions on Automatic Control 19, 716–723.

Archer, L.C., Hutton, S.A., Harman, L., O’Grady, M.N., Kerry, J.P., Poole, W.R., Gargan, P., McGinnity, P., Reed, T.E., 2019. The interplay between extrinsic and intrinsic factors in determining migration decisions in brown trout (Salmo trutta): An experimental study. Frontiers in Ecology and Evolution 7, 222. 10.3389/fevo.2019.00222

Baglinière, J.-L., Hamelet, V., Guéraud, F., Aymes, J.-C., Goulon, C., Richard, A., Josset, Q., Marchand, F., 2020. Guide to the interpretation of the scales and the estimation of the age of brown trout (Salmo trutta) from the French populations., Guides and. ed.

Bal, G., Montorio, L., Rivot, E., Prévost, E., Baglinière, J.-L., Nevoux, M., 2017. Evidence for long-term change in length, mass and migration phenology of anadromous spawners in French Atlantic salmon Salmo salar. Journal of Fish Biology 90, 2375–2393. 10.1111/jfb.13314

Barneche, D.R., Robertson, D.R., White, C.R., Marshall, D.J., 2018. Fish reproductive-energy output increases disproportionately with body size. Science 360, 642–645. 10.1126/science.aao6868

Bartolino, V., Margonski, P., Lindegren, M., Linderholm, H.W., Cardinale, M., Rayner, D., Wennhage, H., Casini, M., 2014. Forecasting fish stock dynamics under climate change: Baltic herring (Clupea harengus) as a case study. Fisheries Oceanography 23, 258–269. 10.1111/fog.12060

Behrenfeld, M.J., O’Malley, R.T., Siegel, D.A., McClain, C.R., Sarmiento, J.L., Feldman, G.C., Milligan, A.J., Falkowski, P.G., Letelier, R.M., Boss, E.S., 2006. Climate-driven trends in contemporary ocean productivity. Nature 444, 752–755. 10.1038/nature05317

Birkeland, K., 1996. Consequences of premature return by sea trout (Salmo trutta) infested with the salmon louse (Lepeophtheirus salmonis Kroyer): migration, growth, and mortality. Canadian Journal of Fisheries and Aquatic Sciences 53, 2808–2813. 10.1139/f96-231

Birkeland, K., Jakobsen, P.J., 1997. Salmon lice, Lepeophtheirus salmonis, infestation as a causal agent of premature return to rivers and estuaries by sea trout, Salmo trutta, juveniles. Environmental Biology of Fishes 49, 129–137. 10.1023/A:1007354632039

Bjørn, P.A., Finstad, B., Kristoffersen, R., 2001. Salmon lice infection of wild sea trout and Arctic char in marine and freshwaters: The effects of salmon farms. Aquaculture Research 32, 947–962. 10.1046/j.1365-2109.2001.00627.x

Blackwell, B.G., Brown, M.L., Willis, D.W., 2000. Relative Weight (Wr) Status and Current Use in Fisheries Assessment and Management. Reviews in Fisheries Science 8, 1–44. 10.1080/10641260091129161

Blann, K., Frost Nerbonne, J., Vondracek, B., 2002. Relationship of Riparian Buffer Type to Water Temperature in the Driftless Area Ecoregion of Minnesota. North American Journal of Fisheries Management 22, 441–451. 10.1577/1548-8675(2002)022<0441:rorbtt>2.0.co;2

Boëns, A., Grellier, P., Lebigre, C., Petitgas, P., 2021. Determinants of growth and selective mortality in anchovy and sardine in the Bay of Biscay. Fisheries Research 239, 105947. 10.1016/j.fishres.2021.105947

Brown, C.J., Fulton, E.A., Hobday, A.J., Matear, R.J., Possingham, H.P., Bulman, C., Christensen, V., Forrest, R.E., Gehrke, P.C., Gribble, N.A., Griffiths, S.P., Lozano-Montes, H., Martin, J.M., Metcalf, S., Okey, T.A., Watson, R., Richardson, A.J., 2010. Effects of climate-driven primary production change on marine food webs: Implications for fisheries and conservation. Global Change Biology 16, 1194–1212. 10.1111/j.1365-2486.2009.02046.x

Burnham, K.P., Anderson, D.R., Burnham, K.P., 2002. Model selection and multimodel inference: a practical information-theoretic approach, 2nd ed. ed. Springer, New York.

Butler, J.R.A., Walker, A.F., 2006. Characteristics of the Sea Trout Salmo trutta (L.) Stock Collapse in the River Ewe (Wester Ross, Scotland), in 1988–2001, in: Harris, G., Milner, N. (Eds.), Sea Trout: Biology, Conservation and Management. Blackwell Science Ltd, pp. 45–59. 10.1002/9780470996027.ch4

Canty, A., Ripley, B., 2021. boot: Bootstrap R (S-Plus) Functions. R package version 1.3–28.

Celtic Sea Trout Project, 2016. Celtic Sea Trout Project - Technical Report to Ireland Wales Territorial Co-operation Programme 2007-2013 (INTERREG 4A. Inland Fisheries Ireland, Dublin.

Crozier, L.G., Hutchings, J. a., 2014. Plastic and evolutionary responses to climate change in fish. Evolutionary Applications 7, 68–87. 10.1111/eva.12135

Davaine, P., Beall, E., Guerri, O., Caraguel, J.M., 1997. Introduction de salmonidés en milieu vierge (îles kerguelen, subantarctique): Enjeux, résultats, perspectives. Bulletin Français de la Pêche et de la Pisciculture 93–110. 10.1051/kmae:1997013

Davidson, I.C., Cove, R.J., Hazlewood, M.S., 2006. Annual Variation in Age Composition, Growth and Abundance of Sea Trout Returning to the River Dee at Chester, 1991– 2003, in: Harris, G., Milner, N. (Eds.), Sea Trout: Biology, Conservation and Management. Proceedings of the First International Sea Trout Symposium, Cardiff, July 2004. Blackwell Publishing Ltd, pp. 76–87.

Davison, A., Hinkley, D., 1997. Bootstrap Methods and Their Applications. Cambridge University Press, Cambridge.

De Eyto, E., Kelly, S., Rogan, G., French, A., Cooney, J., Murphy, M., Nixon, P., Hughes, P., Sweeney, D., McGinnity, P., Dillane, M., Poole, R., 2022. Decadal Trends in the Migration Phenology of Diadromous Fishes Native to the Burrishoole Catchment, Ireland. Front. Ecol. Evol. 10, 915854. 10.3389/fevo.2022.915854

Dȩbowski, P., 2018. The largest Baltic population of sea trout (Salmo trutta L.): Its decline, restoration attempts, and current status. Archives of Polish Fisheries 26, 81–100. 10.2478/aopf-2018-0010

Dodson, J.J., Aubin-Horth, N., Thériault, V., Páez, D.J., 2013. The evolutionary ecology of alternative migratory tactics in salmonid fishes. Biological Reviews 88, 602–625. 10.1111/brv.12019

Dormann, C.F., Elith, J., Bacher, S., Buchmann, C., Carl, G., Carré, G., Marquéz, J.R.G., Gruber, B., Lafourcade, B., Leitão, P.J., Münkemüller, T., Mcclean, C., Osborne, P.E., Reineking, B., Schröder, B., Skidmore, A.K., Zurell, D., Lautenbach, S., 2013. Collinearity: A review of methods to deal with it and a simulation study evaluating their performance. Ecography 36, 27–46. 10.1111/j.1600-0587.2012.07348.x

Eliott, J.M., Eliott, J.A., 2006. A 35-year study of stock-recruitment relationships in a small population of sea trout: assumptions, implications and Limitations for predicting targets., in: Harris, G., Milner, N. (Eds.), Sea Trout: Biology, Conservation and Management. Blackwell Publishing, Oxford, pp. 257–278.

Erkinaro, J., Czorlich, Y., Orell, P., Kuusela, J., Falkegård, M., Länsman, M., Pulkkinen, H., Primmer, C.R., Niemelä, E., 2019. Life history variation across four decades in a diverse population complex of atlantic salmon in a large subarctic river. Canadian Journal of Fisheries and Aquatic Sciences 76, 42–55. 10.1139/cjfas-2017-0343

Euzenat, G., Fournel, F., Fagard, J.L., 2007. Population Dynamics and Stock-Recruitment Relationship of Sea Trout in the River Bresle, Upper Normandy, France. Sea Trout: Biology, Conservation and Management 307–323. 10.1002/9780470996027.ch20

Fagard, J.-L., Beaulaton, L., 2018. Éléments sur l’exploitation par pêche des salmonidés migrateurs en zone côtière et en rivière depuis 1978. Littoral proche de la rivière Bresle, embouchures de cours d’eau de Seine-Maritime. Cours d’eau à salmonidés migrateurs au nord de la Seine. Agence Française pour la Biodiversité.

Ferguson, A., Reed, T.E., Cross, T.F., McGinnity, P., Prodöhl, P.A., 2019. Anadromy, potamodromy and residency in brown trout Salmo trutta: the role of genes and the environment. Journal of Fish Biology. 10.1111/jfb.14005

Fiske, P., Forseth, T., Thorstad, E.B., Bakkestuen, V., Einum, S., Falkegård, M., Garmo, Ø.A., Garseth, Å.H., Skoglund, H., Solberg, M., Utne, K.R., Vollset, K.W., Vøllestad, L.A., Wennevik, V., 2024. Novel large-scale mapping highlights poor state of sea trout populations. Aquatic Conservation: Marine and Freshwater Ecosystems 1–20.

Forget, G., Baglinière, J.L., Marchand, F., Richard, A., Nevoux, M., Durif, C., 2018. A new method to estimate habitat potential for Atlantic salmon (Salmo salar): Predicting the influence of dam removal on the Sélune River (France) as a case study. ICES Journal of Marine Science 75, 2172–2181. 10.1093/icesjms/fsy089

Forseth, T., Barlaup, B.T., Finstad, B., Fiske, P., Gjøsæter, H., Falkegård, M., Hindar, A., Mo, T.A., Rikardsen, A.H., Thorstad, E.B., Vøllestad, L.A., Wennevik, V., 2017. The major threats to Atlantic salmon in Norway. ICES Journal of Marine Science 74, 1496– 1513. 10.1093/icesjms/fsx020

Gargan, P., Kelly, F., Shephard, S., Whelan, K., 2016. Temporal variation in sea trout Salmo trutta life history traits in the Erriff River, western Ireland. Aquaculture Environment Interactions 8, 675–689. 10.3354/aei00211

Gargan, P.G., Poole, W.R., Forde, G.P., 2006. A review of the status of irish sea trout stocks., in: Harris, G., Milner, N. (Eds.), Sea Trout: Biology, Conservation and Management. Blackwell Science Ltd, pp. 25–44. 10.1002/9780470996027.ch3

Gohin, F., Van der Zande, D., Tilstone, G., Eleveld, M.A., Lefebvre, A., Andrieux-Loyer, F., Blauw, A.N., Bryère, P., Devreker, D., Garnesson, P., Hernández Fariñas, T., Lamaury, Y., Lampert, L., Lavigne, H., Menet-Nedelec, F., Pardo, S., Saulquin, B., 2019. Twenty years of satellite and in situ observations of surface chlorophyll-a from the northern Bay of Biscay to the eastern English Channel. Is the water quality improving? Remote Sensing of Environment 233, 111343. 10.1016/j.rse.2019.111343

Gross, M.R., Coleman, R.M., McDowall, R.M., 1988. Aquatic productivity and the evolution of diadromous fish migration. Science 239, 1291–1293. 10.1126/science.239.4845.1291

Hard, J.J., Gross, M.R., Heino, M., Hilborn, R., Kope, R.G., Law, R., Reynolds, J.D., 2008. Evolutionary consequences of fishing and their implications for salmon. Evolutionary Applications 1, 388–408. 10.1111/j.1752-4571.2008.00020.x

Helle, J., Martinson, E., Eggers, D., 2007. Influence of salmon abundance and ocean conditions on body size of Pacific salmon. N. Pac. Anadr. Fish 289–298.

Hixon, M. a, Johnson, D.W., Sogard, S.M., 2014. BOFFFFs: on the importance of conserving old-growth age structure in fishery populations. ICES Journal of Marine Science 71, 2171–2185. 10.1093/icesjms/fst200

Hoegh-Guldberg, O., Bruno, J.F., 2010. The impact of climate change on the world’s marine ecosystems. Science 328, 1523–1528. 10.1126/science.1189930

Hughes, B.B., Beas-Luna, R., Barner, A.K., Brewitt, K., Brumbaugh, D.R., Cerny-Chipman, E.B., Close, S.L., Coblentz, K.E., De Nesnera, K.L., Drobnitch, S.T., Figurski, J.D., Focht, B., Friedman, M., Freiwald, J., Heady, K.K., Heady, W.N., Hettinger, A., Johnson, A., Karr, K.A., Mahoney, B., Moritsch, M.M., Osterback, A.M.K., Reimer, J., Robinson, J., Rohrer, T., Rose, J.M., Sabal, M., Segui, L.M., Shen, C., Sullivan, J., Zuercher, R., Raimondi, P.T., Menge, B.A., Grorud-Colvert, K., Novak, M., Carr, M.H., 2017. Long-Term studies contribute disproportionately to ecology and policy. BioScience 67, 271–278. 10.1093/biosci/biw185

Hutchings, J. a, Jones, M.E.B., 1998. Life history variation and growth rate thresholds for maturity in Atlantic salmon, Salmo salar. Canadian Journal of Fisheries and Aquatic Sciences 55 (Suppl., 22–47. 10.1139/cjfas-55-S1-22

ICES, 2020. Working Group with the Aim to Develop Assessment Models and Establish Biological Refer-ence Points for Sea Trout (Anadromous Salmo trutta) Populations (WGTRUTTA; outputs from 2019 meeting). ICES Scientific Reports. 10.17895/ices.pub.7431

Ishida, Y., Ito, S., Kaeriyama, M., McKinnell, S., Nagasawa, K., 1993. Recent changes in age and size of chum salmon (Oncorhynchus keta) in the North Pacific Ocean and possible causes. Canadian Journal of Fisheries and Aquatic Sciences 50, 290–295. 10.1139/f93-033

Järvi, T.H., 1940. Sea trout in the Bothnian Bay. Acta Zoologica Fennica 1–29.

Jeffrey, K.M., Côté, I.M., Irvine, J.R., Reynolds, J.D., 2017. Changes in body size of Canadian Pacific salmon over six decades. Canadian Journal of Fisheries and Aquatic Sciences 74, 191–201. 10.1139/cjfas-2015-0600

Jonsson, B., L’Abée-Lund, J.H., 1993. Latitudinal clines in life-history variables of anadromous brown trout in Europe. Journal of Fish Biology 43, 1–16. 10.1111/j.1095-8649.1993.tb01175.x

Josset, Q., Beaulaton, L., Romakkaniemi, A., Nevoux, M., 2023. Data for “Changes in length-at-first return of a sea trout (Salmo trutta) population in northern France.” 10.57745/FPHLBT

Kendall, N.W., Quinn, T.P., 2011. Length and age trends of Chinook salmon in the Nushagak River, Alaska, related to commercial and recreational fishery selection and exploitation. Transactions of the American Fisheries Society 140, 611–622. 10.1080/00028487.2011.585575

Kennedy, R.J., Crozier, W.W., 2010. Evidence of changing migratory patterns of wild Atlantic salmon Salmo salar smolts in the River Bush, Northern Ireland, and possible associations with climate change. Journal of Fish Biology 76, 1786–1805. 10.1111/j.1095-8649.2010.02617.x

Knutsen, J.A., Knutsen, H., Gjøsæter, J., Jonsson, B., 2001. Food of anadromous brown trout at sea. Journal of Fish Biology 59, 533–543. 10.1006/jfbi.2001.1662

Knutsen, J.A., Knutsen, H., Olsen, E.M., Jonsson, B., 2004. Marine feeding of anadromous *Salmo trutta* during winter. Journal of Fish Biology 64, 89–99. 10.1046/j.1095-8649.2003.00285.x

Kozlowski, J., 2006. Why life-histories are diverse. Polish Journal of Ecology 54, 585–605.

L’Abee-Lund, J.H., Jonsson, B., Jensen, A.J., Saettem, L.M., Heggberget, T.G., Johnsen, B.O., Naesje, T.F., 1989. Latitudinal Variation in Life-History Characteristics of Sea-Run Migrant Brown Trout Salmo trutta. The Journal of Animal Ecology 58, 525. 10.2307/4846

Law, R., 2000. Fishing, selection, and phenotypic evolution. ICES Journal of Marine Science 57, 659–668. 10.1006/jmsc.2000.0731

Legrand, M., Briand, C., Buisson, L., Besse, T., Artur, G., Azam, D., Baisez, A., Barracou, D., Bourré, N., Carry, L., Caudal, A.L., Corre, J., Croguennec, E., Der Mikaélian, S., Josset, Q., Le Gurun, L., Schaeffer, F., Toussaint, R., Laffaille, P., 2021. Diadromous fish modified timing of upstream migration over the last 30 years in France. Freshwater Biology 66, 286–302. 10.1111/fwb.13638

Lewis, B., Grant, W.S., Brenner, R.E., Hamazaki, T., 2015. Changes in size and age of chinook salmon oncorhynchus tshawytscha returning to Alaska. PLoS ONE 10, 1–17. 10.1371/journal.pone.0130184

Maisse, G., Mourot, B., Breton, B., Fostier, A., Marcuzzi, O., Le Bail, P.Y., Baglinière, J.L., Richard, A., 1991. Sexual maturity in sea trout, Salmo trutta L., running up the river Calonne (Normandy, France) at the “finnock” stage. Journal of Fish Biology 39, 705– 715. 10.1111/j.1095-8649.1991.tb04400.x

Marchand, F., Aymes, J.-C., Gueraud, F., Guillard, J., Goulon, C., Hamelet, V., Lange, F., Prévost, E., Baglinière, J.L., Beaulaton, L., Penil, C., Azam, D., 2019. Colisa, the collection of ichtyological samples. 10.15454/D3ODJM

Marshall, D.J., White, C.R., 2019. Have We Outgrown the Existing Models of Growth? Trends in Ecology & Evolution 34, 102–111. 10.1016/j.tree.2018.10.005

McCullagh, P., Nelder, J.A., 1989. Generalized Linear Models, Second Edition., Chapman&Hall. ed, Monographs on Statistics and Applied Probability.

McKinney, G.J., Hale, M.C., Goetz, G., Gribskov, M., Thrower, F.P., Nichols, K.M., 2015. Ontogenetic changes in embryonic and brain gene expression in progeny produced from migratory and resident Oncorhynchus mykiss. Molecular Ecology 24, 1792– 1809. 10.1111/mec.13143

Milner, N., Potter, E., Roche, W., Tysklin, N., Davidson, I.C., King, J., Coyne, J., Davies, C., 2017. Variation in sea trout (Salmo trutta) abundance and life histories in the Irish Sea., in: Graeme Harris. Ed. (Ed.), Sea Trout: Science & Management. Proceedings of the 2nd International Sea Trout Symposium, October 2015, Dundalk, Ireland.

Möllmann, C., Diekmann, R., 2012. Marine Ecosystem Regime Shifts Induced by Climate and Overfishing. A Review for the Northern Hemisphere, Advances in Ecological Research. 10.1016/B978-0-12-398315-2.00004-1

Morita, K., Fukuwaka, M.A., 2007. Why age and size at maturity have changed in Pacific salmon. Marine Ecology Progress Series 335, 289–294. 10.3354/meps335289

Neuheimer, A.B., Taggart, C.T., 2010. Can changes in length-at-age and maturation timing in Scotian Shelf haddock (Melanogrammus aeglefinus) be explained by fishing? Canadian Journal of Fisheries and Aquatic Sciences 67, 854–865. 10.1139/F10-025

Nevoux, M., Finstad, B., Davidsen, J.G., Finlay, R., Josset, Q., Poole, R., Höjesjö, J., Aarestrup, K., Persson, L., Tolvanen, O., Jonsson, B., 2019. Environmental influences on life history strategies in partially anadromous brown trout (Salmo trutta, Salmonidae). Fish and Fisheries 1–32. 10.1111/faf.12396

Niiranen, S., Yletyinen, J., Tomczak, M.T., Blenckner, T., Hjerne, O., Mackenzie, B.R., Müller-Karulis, B., Neumann, T., Meier, H.E.M., 2013. Combined effects of global climate change and regional ecosystem drivers on an exploited marine food web. Global Change Biology 19, 3327–3342. 10.1111/gcb.12309

Ohlberger, J., Schindler, D.E., Brown, R.J., Harding, J.M.S., Adkison, M.D., Munro, A.R., Horstmann, L., Spaeder, J., 2020. The reproductive value of large females: Consequences of shifts in demographic structure for population reproductive potential in Chinook Salmon. Canadian Journal of Fisheries and Aquatic Sciences 77, 1292–1301. 10.1139/cjfas-2020-0012

Ohlberger, J., Schindler, D.E., Ward, E.J., Walsworth, T.E., Essington, T.E., 2019. Resurgence of an apex marine predator and the decline in prey body size. Proceedings of the National Academy of Sciences of the United States of America 116, 26682–26689. 10.1073/pnas.1910930116

Ohlberger, J., Ward, E.J., Schindler, D.E., Lewis, B., 2018. Demographic changes in Chinook salmon across the Northeast Pacific Ocean. Fish and Fisheries 19, 533–546. 10.1111/faf.12272

Ojuok, J.E., Njiru, M., Ntiba, M.J., Mavuti, K.M., 2007. The effect of overfishing on the life-history strategies of Nile tilapia, Oreochromis niloticus (L.) in the Nyanza Gulf of Lake Victoria, Kenya. Aquatic Ecosystem Health and Management 10, 443–448. 10.1080/14634980701708107

Oke, K.B., Cunningham, C.J., Westley, P.A.H., Baskett, M.L., Carlson, S.M., Clark, J., Hendry, A.P., Karatayev, V.A., Kendall, N.W., Kibele, J., Kindsvater, H.K., Kobayashi, K.M., Lewis, B., Munch, S., Reynolds, J.D., Vick, G.K., Palkovacs, E.P., 2020. Recent declines in salmon body size impact ecosystems and fisheries. Nature Communications 11, 1–13. 10.1038/s41467-020-17726-z

Perez, K.O., Munch, S.B., 2010. Extreme selection on size in the early lives of fish. Evolution 64, 2450–2457. 10.1111/j.1558-5646.2010.00994.x

Perrier, C., Guyomard, R., Bagliniere, J.L., Nikolic, N., Evanno, G., 2013. Changes in the genetic structure of Atlantic salmon populations over four decades reveal substantial impacts of stocking and potential resiliency. Ecology and Evolution 3, 2334–2349. 10.1002/ece3.629

Quinn, T.P., McGinnity, P., Reed, T.E., 2016. The paradox of “premature migration” by adult anadromous salmonid fishes: Patterns and hypotheses. Canadian Journal of Fisheries and Aquatic Sciences 73, 1015–1030. 10.1139/cjfas-2015-0345

R Core Team, 2020. R: A language and environment for statistical computing.

Reed, T.E., Prodöhl, P., Bradley, C., Gilbey, J., McGinnity, P., Primmer, C.R., Bacon, P.J., 2019. Heritability estimation via molecular pedigree reconstruction in a wild fish population reveals substantial evolutionary potential for sea age at maturity, but not size within age classes. Can. J. Fish. Aquat. Sci. 76, 790–805. 10.1139/cjfas-2018-0123

Reznick, D.A., Bryga, H., Endler, J.A., 1990. Experimentally induced life-history evolution in a natural population. Nature 346, 357–359. 10.1038/346357a0

Richard, A., 1981. Observations préliminaires sur les populations de truite de mer (Salmo trutta L.) en Basse-Normandie. Bulletin Français de la Pêche et de la Pisciculture 283, 114–124.

Rikardsen, a. H., Amundsen, P. a., Knudsen, R., Sandring, S., 2006. Seasonal marine feeding and body condition of sea trout (Salmo trutta) at its northern distribution. ICES Journal of Marine Science 63, 466–475. 10.1016/j.icesjms.2005.07.013

Rikardsen, A.H., Dempson, J.B., Amundsen, P.-A., Bjorn, P.A., Finstad, B., Jensen, A.J., 2007. Temporal variability in marine feeding of sympatric Arctic charr and sea trout. Journal of Fish Biology 70, 837–852. 10.1111/j.1095-8649.2007.01345.x

Russell, I.C., Aprahamian, M.W., Barry, J., Davidson, I.C., Fiske, P., Ibbotson, A.T., Kennedy, R.J., Maclean, J.C., Moore, A., Otero, J., Potter, T. (E. C.E.), Todd, C.D., 2012. The influence of the freshwater environment and the biological characteristics of Atlantic salmon smolts on their subsequent marine survival. ICES Journal of Marine Science 69, 1563–1573. 10.1093/icesjms/fsr208

Satterthwaite, W.H., Mohr, M.S., O’Farrell, M.R., Wells, B.K., 2012. A Bayesian hierarchical model of size-at-age in ocean-harvested stocks-quantifying effects of climate and temporal variability. Canadian Journal of Fisheries and Aquatic Sciences 69, 942–954. 10.1139/F2012-036

Serra-Llinares, R.M., Bøhn, T., Karlsen Nilsen, R., Freitas, C., Albretsen, J., Haraldstad, T., Thorstad, E.B., Elvik, K.M.S., Bjørn, P.A., 2020. Impacts of salmon lice on mortality, marine migration distance and premature return in sea trout. Marine Ecology Progress Series 635, 151–168. 10.3354/MEPS13199

Spitz, J., Dupuis, L., Becquet, V., Dubief, B., Trites, A.W., 2015. Diet of the harbour seal Phoca vitulina: Implication for the flatfish nursery in the Bay of Somme (English Channel, France). Aquatic Living Resources 28, 11–19. 10.1051/alr/2015001

Stewart, J., 2011. Evidence of age-class truncation in some exploited marine fish populations in New South Wales, Australia. Fisheries Research 108, 209–213. 10.1016/j.fishres.2010.11.017

Suuronen, P., Lehtonen, E., 2012. The role of salmonids in the diet of grey and ringed seals in the Bothnian Bay, northern Baltic Sea. Fisheries Research 125–126, 283–288. 10.1016/j.fishres.2012.03.007

Tattam, I.A., Ruzycki, J.R., McCormick, J.L., Carmichael, R.W., 2015. Length and Condition of Wild Chinook Salmon Smolts Influence Age at Maturity. Transactions of the American Fisheries Society 144, 1237–1248. 10.1080/00028487.2015.1082503

Thomas, A.C., Nelson, B.W., Lance, M.M., Deagle, B.E., Trites, A.W., 2017. Harbour seals target juvenile salmon of conservation concern. Canadian Journal of Fisheries and Aquatic Sciences 74, 907–921. 10.1139/cjfas-2015-0558

Thomsen, D.S., Koed, A., Nielsen, C., Madsen, S.S., 2007. Overwintering of sea trout (Salmo trutta) in freshwater: Escaping salt and low temperature or an alternate life strategy? Canadian Journal of Fisheries and Aquatic Sciences 64, 793–802. 10.1139/F07-059

Thorstad, E.B., Todd, C.D., Uglem, I., Bjørn, P.A., Gargan, P.G., Vollset, K.W., Halttunen, E., Kålås, S., Berg, M., Finstad, B., 2016. Marine life of the sea trout. Marine Biology 163, 1–19. 10.1007/s00227-016-2820-3

Thorstad, E.B., Todd, C.D., Uglem, I., Bjørn, P.A., Gargan, P.G., Vollset, K.W., Halttunen, E., Kålås, S., Berg, M., Finstad, B., 2015. Effects of salmon lice Lepeophtheirus salmonis on wild sea trout salmo trutta - A literature review. Aquaculture Environment Interactions 7, 91–113. 10.3354/aei00142

Tréhin, C., Rivot, E., Lamireau, L., Meslier, L., Besnard, A., Gregory Stephen, D., Nevoux, M., 2021. Growth during the first summer at sea modulates sex-specific maturation schedule in Atlantic salmon. Canadian Journal of Fisheries and Aquatic Sciences 1–40. 10.1139/cjfas-2020-0236

Waco, K.E., Taylor, W.W., 2010. The influence of groundwater withdrawal and land use changes on brook charr (Salvelinus fontinalis) thermal habitat in two coldwater tributaries in Michigan, U.S.A. Hydrobiologia 650, 101–116. 10.1007/s10750-010-0204-0

Walsh, M.R., Reznick, D.N., 2011. Experimentally induced life-history evolution in a killifish in response to the introduction of guppies. Evolution 65, 1021–1036. 10.1111/j.1558-5646.2010.01188.x

Witten, P.E., Hall, B.K., 2001. The salmon kype. How does it grow? What’s its purpose? Does it disappear after spawning? Atlantic Salmon Journal 36–39.

